# Reverse engineering the anti-MUC1 hybridoma antibody 139H2 by mass spectrometry-based *de novo* sequencing

**DOI:** 10.1101/2023.07.05.547778

**Authors:** Weiwei Peng, Koen C.A.P. Giesbers, Marta Šiborová, J. Wouter Beugelink, Matti F. Pronker, Douwe Schulte, John Hilkens, Bert J.C. Janssen, Karin Strijbis, Joost Snijder

## Abstract

Mucin 1 (MUC1) is a transmembrane mucin expressed at the apical surface of epithelial cells at different mucosal surfaces including breast and intestine. In the gastrointestinal tract, MUC1 has a barrier function against bacterial invasion, but can also serve as an entry receptor for pathogenic *Salmonella* bacteria. Moreover, MUC1 is well known for its aberrant expression and glycosylation in adenocarcinomas The MUC1 extracellular domain contains a variable number of tandem repeats (VNTR) of 20 amino acids, which are heavily *O*-linked glycosylated.. Monoclonal antibodies against the MUC1 VNTR can be powerful tools because of their multiplicity of binding and possible applications in the diagnosis and treatment of MUC1-expressing cancers. One such antibody is the hybridoma mouse monoclonal 139H2, which is also widely used as a research tool to study non-cancer MUC1. Here we report direct mass spectrometry-based sequencing of hybridoma-derived 139H2 IgG, which enabled reverse engineering of a recombinant 139H2. The performance of the reverse engineered 139H2 IgG and its Fab fragment were validated by comparison to the hybridoma-derived product in Western blot and immunofluorescence microscopy. The reverse engineering of 139H2 allowed us to characterize binding to the VNTR peptide epitope by surface plasmon resonance (SPR) and solve the crystal structure of the 139H2 Fab fragment in complex with the MUC1 VNTR peptide. These analyses reveal the molecular basis for 139H2 binding specificity to MUC1 and its tolerance to *O*-glycosylation of the VNTR. The available sequence of 139H2 will allow further development of MUC1-related diagnostics, targeting and treatment strategies.

## Introduction

The mucin MUC1 is a transmembrane glycoprotein expressed by epithelial cells at different mucosal surfaces including breast tissue, the airways and gastrointestinal tract. The full-length MUC1 protein extends 200-500 nm from the apical surface of epithelial cells and is therefore an important component of the glycocalyx^1,2^. At the mucosal surface, MUC1 has an essential barrier function against bacterial and viral invasion^3,4^ but it can also be used as entry receptor by pathogenic *Salmonella* species ^5^. Using knockout mice, it was demonstrated that MUC1 has anti-inflammatory functions^6–8^. However, MUC1 is most well-known for its aberrant expression and glycosylation in different types of adenocarcinomas

The full-length MUC1 heterodimer consists of an extracellular domain with a variable number of tandem repeats (VNTR) of 20 amino acids, which are heavily *O*-linked glycosylated, a non-covalently attached SEA domain, a transmembrane domain, and a cytoplasmic tail with signaling capacity (see Figure 1). The VNTR region consists of repeats of 20 amino acids with the sequence GSTAPPAHGVTSAPDTRPAP^10,11^. Each repeat contains five serine and threonine residues that can be *O*-linked glycosylated and experiments with synthetic MUC1 fragments demonstrated a high glycosylation occupancy at these residues^12^. In healthy tissue, the *O*-glycans on the MUC1 VNTR predominantly consist of elongated core 2 structures, while it remains restricted to predominant core 1 structures in many cancerous cells^13,14^.

**Figure 1.**
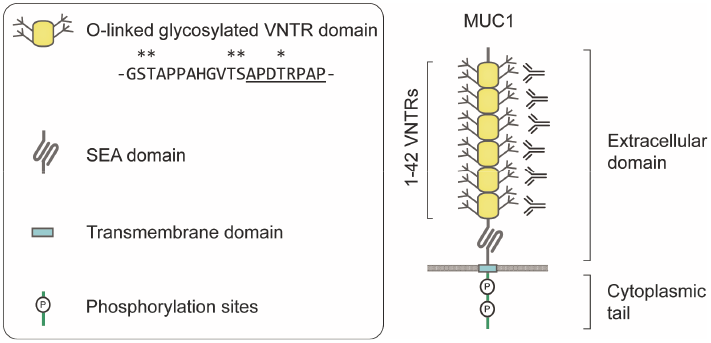
Schematic overview of MUC1 domain structure. VNTR: Variable Number of Tandem Repeats. SEA: domain name from initial identification in a <u>s</u>perm protein, <u>e</u>nterokinase, and <u>a</u>grin.

The overexpression and altered glycosylation of MUC1 in cancerous cells makes it a potentially viable candidate target for cancer immunotherapy. In addition, MUC1 could be an interesting target for therapeutic strategies that require delivery to the (healthy) mucosal surface. Monoclonal antibodies against the MUC1 VNTR can be powerful tools because of their multiplicity of binding and possible applications in the diagnosis and treatment of MUC1-expressing cancers. Since the late 1980’s, several monoclonal antibodies against MUC1 have been described and explored for the diagnosis and treatment of MUC1 overexpressing cancers^15,16^. Peptide mapping experiments have revealed that many such monoclonal antibodies target a similar region within the VNTR of MUC1, resulting in the definition of an immunodominant peptide corresponding to the subsequence APDTRPAP^17^. One such antibody is 139H2, a hybridoma monoclonal antibody that was raised against human breast cancer plasma membranes^15,16^. In different studies, 139H2 has been applied for the diagnostics of MUC1-overexpressing cancers and radioimmunotherapy^15,16,18^. In addition, the antibody is also widely applied as a research tool in Western blot, ELISA, immunohistochemistry and immunofluorescence microscopy to study MUC1 biology^16,19,20^. To make this antibody available for general use, we set out to determine its sequence based on the available hybridoma-derived product. Recently we have reported a method to reverse engineer monoclonal antibodies by determining the sequence directly from the purified protein product based on liquid chromatography coupled to mass spectrometry (LC-MS), using a bottom-up proteomics approach^21–24^. Here we applied this method to obtain the full sequence of 139H2. The sequence was successfully validated by comparing the performance of the reverse engineered 139H2 and its Fab fragment to the hybridoma-derived product in Western blot and immunofluorescence microscopy. Reverse engineering 139H2 enabled us to characterize binding to the immunodominant peptide epitope within the MUC1 VNTR by surface plasmon resonance (SPR) and map out the epitope by solving a crystal structure of the 139H2 Fab fragment in complex with the APDTRPAP peptide. These analyses reveal the molecular basis for 139H2 binding to MUC1 and illustrate a remarkable diversity of binding modes to the immunodominant epitope in comparison to other reported structures of anti-MUC1 monoclonals targeting the VNTR.

## Result

### De novo sequencing by bottom-up mass spectrometry

The goal of our study was to obtain the sequence of the full length 139H2 IgG antibody using a bottom-up proteomics approach. As a starting point, we used 139H2 IgG hybridoma supernatant and purified the antibody using protein G affinity resin. The purified IgG was digested with a panel of 4 proteases in parallel (trypsin, chymotrypsin, α-lytic protease, and thermolysin) to generate overlapping peptides for the LC-MS/MS analysis, using a hybrid fragmentation scheme with stepped high-energy collision dissociation (sHCD) and electron-transfer high energy collision dissociation (EThcD) on all peptide precursors. The peptide sequences were predicted from the MS/MS spectra using PEAKS and assembled into the full-length heavy and light chain sequences using the in-house developed software Stitch. This resulted in the identification of a mouse IgG1 antibody with an IGHV1-53 heavy chain paired with an IGKV8-30 light chain (the full sequence is provided in the Supplementary Information). The depth of coverage for the complementarity determining regions (CDRs) varies from around 10 to 100, indicating a high sequence accuracy (see Supplementary Figure S1). Examples of MS/MS spectra supporting the CDRs of both heavy chain and light chain are shown in Figure 2. Comparison to the inferred germline precursors indicate a typical moderate level of somatic hypermutation (3% in the light chain; 10% in the heavy chain), with some notable mutations in the framework regions, also directly flanking CDRH2.

**Figure 2.**
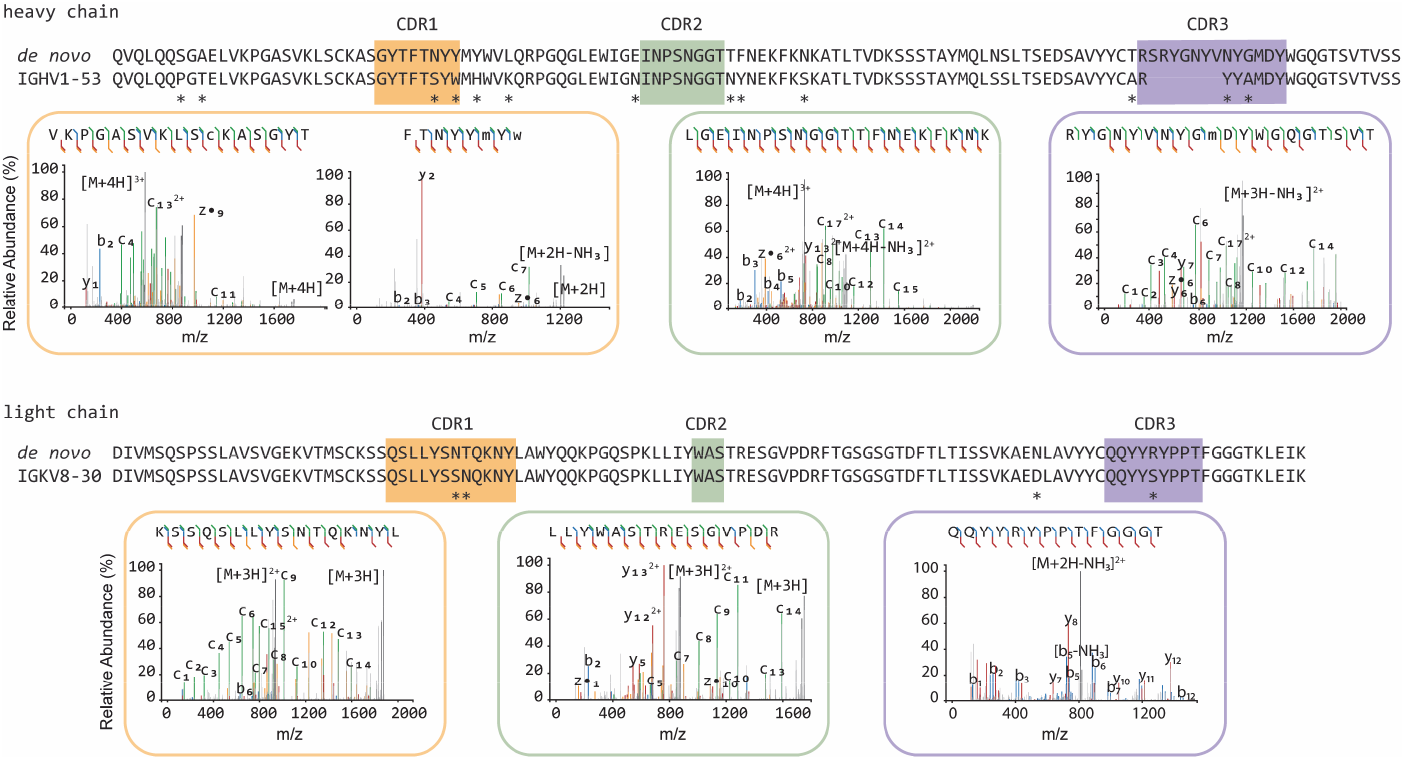
De novo sequencing of the hybridoma 139H2 based on bottom-up proteomics. The variable region alignment to the inferred germline sequence is shown for both heavy and light chains. Positions with putative somatic hypermutation are highlighted with asterisks (*). The MS/MS spectra supporting the CDR regions are shown beneath the sequence alignment, b/y ions are indicated in blue and red, while c/z ions are indicated in green and yellow.

**Figure 3.**
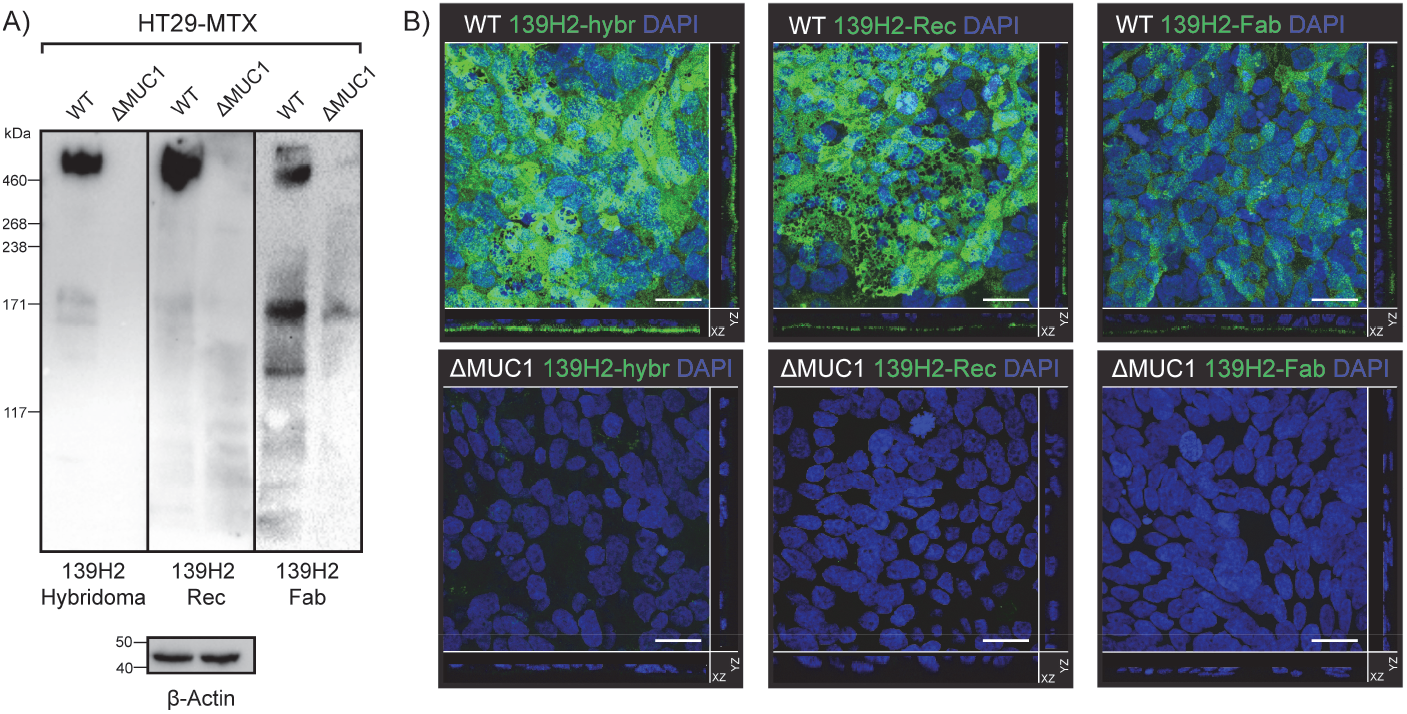
Validation of synthetic recombinant 139H2 following the mass spectrometry-derived sequence. (A) Immunoblot analysis of lysates of intestinal epithelial HT29-MTX and HT29-MTX ΔMUC1 cells with the original hybridoma-derived 139H2 IgG antibody and synthetic recombinant 139H2. (B) Immunofluorescence confocal microscopy imaging of confluent HT29-MTX and HT29-MTX ΔMUC1 monolayers. Cells were stained for nuclei (DAPI, blue) and MUC1 (139H2, green). The signal of the 139H2 Fab was enhanced to compensate for the expected low signal/binding. White scale bars represent 20 μm.

### Validation of the experimentally determined 139H2 sequence

The experimentally determined sequences of the 139H2 variable domains were codon optimized for mammalian expression and subcloned into expression vectors with the mouse IgG1 heavy chain (with an 8xHis-tag) and the kappa light chain backbones (see Supplementary Information for the full amino acid sequences). Co-transfection of the two plasmids in HEK293 cells yielded ca. 10 mg from a 1 L culture following His-trap purification (see Supplementary Figure S2). Additionally, the Fragment antigen-binding (Fab) region was expressed to study the monovalent binding to MUC1. The recombinant 139H2 and Fab were then compared with the hybridoma-derived 139H2 in Western blot and confocal immunofluorescence microscopy.

To investigate the specificity of the recombinant 139H2 antibody for MUC1, we performed immunoblot analysis on lysates of the methotrexate-adapted human colon cancer cell line HT29-MTX, known for its high MUC1 expression, and a MUC1 knockout of the same cell line that was previously described (see Figure 2)^5^. The original hybridoma-derived 139H2 recognizes one predominant band at an estimated molecular weight of 600 kDa, corresponding to full length MUC1, and this band is absent in lysates of the MUC1-knockout cells. The recombinant 139H2 showed the same binding pattern. In confocal immunofluorescence microscopy, original hybridoma-derived 139H2 stains MUC1 at the apical surface in a confluent culture of HT29-MTX, and this signal is reduced to background in the MUC1-knockout cell line. A similar staining is observed with the recombinant 139H2. Western blot and immunofluorescence microscopy using the monovalent Fab fragment also showed specific binding to MUC1 in the wild type background but with reduced avidity compared to the full bivalent IgG molecule. These results confirm that the reverse engineered 139H2 antibody is functional and recognizes the full length MUC1 glycoprotein at the apical surface of intestinal epithelial cells.

### Epitope mapping of 139H2

Using the reverse engineered 139H2 product, we next characterized binding to the immunodominant epitope APDTRPAPG within the MUC1 VNTR. Binding to the synthetic peptide, including an N-terminal biotin and short peptide linker for immobilization to the SPR substrate (*i*.*e*. biotin-GGS-APDTRPAPG), was determined by SPR. Binding of the full IgG was characterized by a high and low affinity phase with dissociation constants of 17×10^-9^ M and 43×10^-7^ M, respectively (Figure S3). We interpret this biphasic binding as an avidity-enhanced bivalent mode (both Fab arms engaged with epitope, high affinity), and a monovalent mode (single Fab arm, low affinity) of binding, respectively. In line with this interpretation, binding to a recombinant monovalent 139H2 Fab yielded a dissociation constant of 45×10^-7^ M, similar to the low affinity binding phase of the full IgG.

To better understand the molecular basis of 139H2 binding to the immunodominant epitope within the VNTR we determined a crystal structure of the Fab fragment in complex with the synthetic APDTRPAPG peptide (without N-terminal biotin or peptide linker). Crystals diffracted to a resolution of 2.5 Å and a structure was solved using molecular replacement with a ColabFold model of the 139H2 Fab. This also revealed clear density for the peptide epitope in contact with the CDRs of 139H2 (see Supplementary Table S1 and Supplementary Figure S4).

The APDTRPAPG peptide binds diagonally across the cleft between the heavy and light chains, making direct contact with all CDRs, except CDRL2 (see Figure 4 and Supplementary Table S2). Contact points between the peptide and the 139H2 Fab include hydrogen bonds with the peptide backbone at 6 out of 8 positions. Both the aspartic acid and arginine residues within the epitope make salt bridges with side chains from 139H2. While D3 interacts with R99 within CDRL1, R5 interacts with E50 and T59 near CDRH2, in addition to a stacking interaction with Y100 in CDRL3. Neither residue E50 nor T59 in 139H2 is formally part of CDRH2, though both residues directly flank the loop. Previous studies on the binding specificity of 139H2 have shown that R5 of the epitope is crucial for 139H2 binding. The crystal structure reported here shows that interactions with R5 are mediated by residues in 139H2 that are formally part of the framework regions of the heavy chain, but both mutated compared to the inferred germline precursors (see Figure 2). Two additional framework mutations in the heavy chain, *i*.*e*. Y35 and T97, appear indirectly involved in MUC1 binding by positioning CDRH3 through hydrogen bonds with N106 and the backbone of Y111, respectively (see Supplementary Figure S5). Finally, the T4 residue of the APDTRPAPG epitope is a known glycosylation site, although 139H2 binding is reported to be unaffected by the presence of a single *O*-linked GalNAc at this position^14,25^. The crystal structure reported here shows the T4 side chain to be pointing outwards from the 139H2 paratope with no indication of potential clashes that would preclude binding of the epitope with glycosylated APDTRPAPG at the T4 position. In line with this previous report and our own structural data, we also found that 139H2 binds equally well to MUC1 reporter constructs with different types of *O*-linked glycans (Supplementary Figure S6).

**Figure 4.**
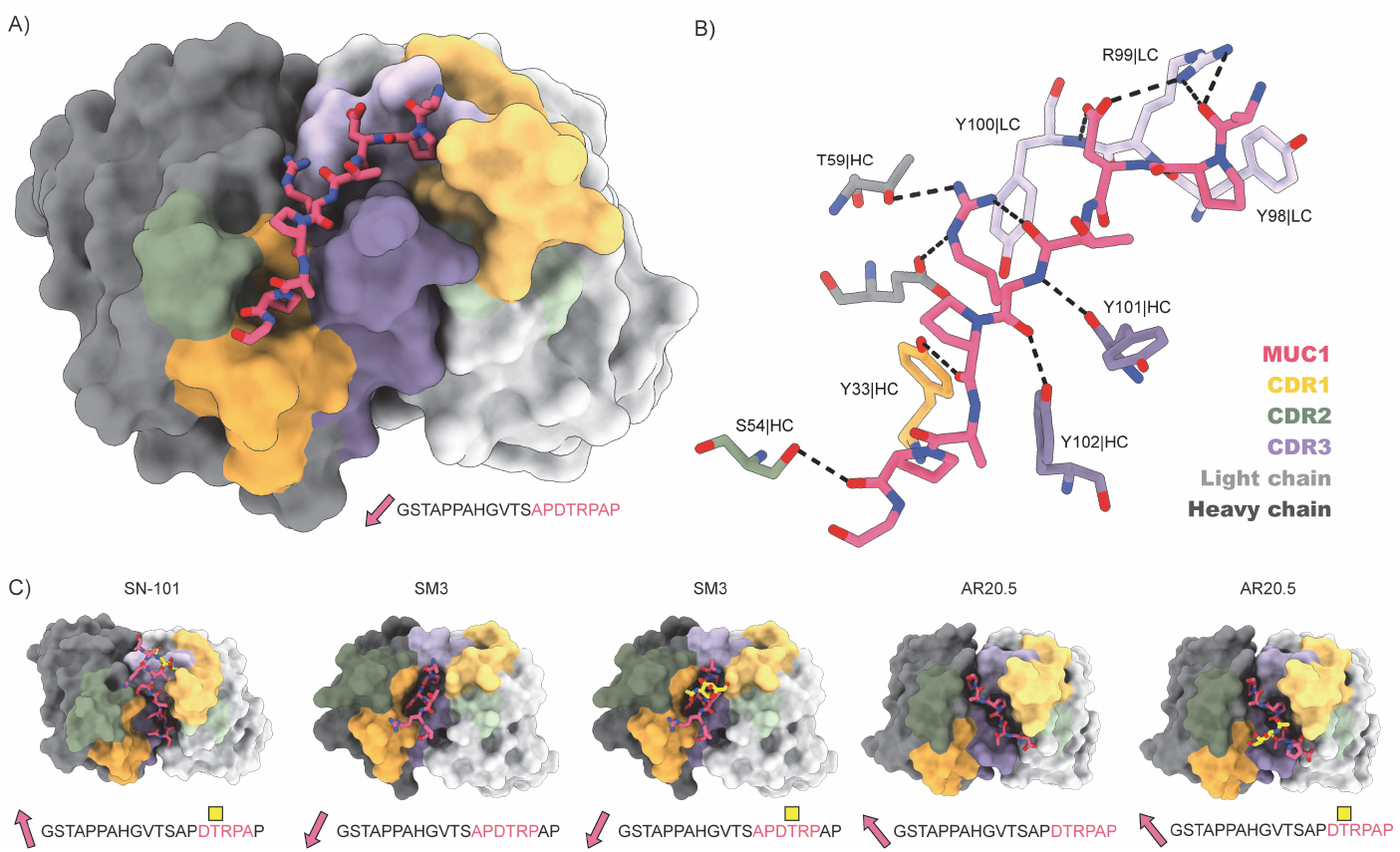
Structure of 139H2 Fab in complex with MUC1 peptide. (A) Surface representation of the Fab with CDRs highlighted in colours and MUC1 peptide shown as a model. N-to C-terminus direction of MUC1 peptides is shown as a pink arrow. (B) Interactions of interactions between 139H2 Fab and MUC1 peptide. (C) Comparison with previously reported structures of monoclonal anti-MUC1 antibodies targeting the VNTR. Glycosylated residues of the epitope are depicted by yellow square above.

Comparison with previously reported structures of monoclonal anti-MUC1 antibodies targeting the VNTR reveal a striking diversity in the modes of binding (a full overview of reported structures is listed in Supplementary Table S3)^26–36^. Monoclonal antibodies 14A, 16A, and 5E5 all target a different region within the VNTR. While monoclonal antibodies SM3, SN101, and AR20.5 all bind to the same immunodominant epitope of the VNTR as 139H2, the peptide is either shifted or oriented in the opposite direction relative to the cleft between the heavy and light chains. For SN101 and AR20.5, the peptide runs across this cleft in the opposite direction compared to 139H2. In SM3 the peptide is oriented in a similar direction but shifted by approximately 2 residues such that both D3 and R5 are contacting different CDRs. In contrast to 139H2, each of the monoclonals compared above bind stronger to the glycosylated epitope. In the case of AR20.5 and SN101 this specificity can be explained by direct contacts made between the glycan and CDRs of the antibody. However, for SM3 the orientation of the glycosylated T4 residue is more similar to 139H2. In SM3 the GalNAc residue makes an additional hydrogen bond with a tyrosine in CDRL1. A similar interaction is predicted for 139H2, albeit through a different group of the GalNAc residue (see Supplementary Figure S7).

## Discussion

Our study demonstrates how direct mass spectrometry-based protein sequencing enables the reconstruction of antibodies from hybridoma supernatants. In addition to recovering such precious resources for research and therapeutic applications, it also contributes to open and reproducible science by making the sequences of crucial monoclonal antibody reagents more readily available and accessible. Poorly defined (monoclonal) antibody products have notoriously been a challenge to reproducibility in life science research and the present work shows that MS-based sequencing can offer helpful improvements in this regard^37,38^.

The reverse-engineered anti-MUC1 monoclonal antibody 139H2 reported here is suitable for Western blotting and immunofluorescence microscopy and is likely suitable for other applications in FACS sorting of MUC1 positive cells, immunohistochemistry and ELISA, as demonstrated for the original hybridoma-derived product^16,19,20^. We show that 139H2 binds the immunodominant epitope of the VNTR in a unique way compared to previously described monoclonal antibodies against MUC1. Because of its previously reported glycan-independent binding, which we supported in this study by the determined structure in complex with the epitope, the 139H2 antibody is an important tool for current and future MUC1 research.

## Methods

### Purification of 139H2 from hybridoma cultures supernatant

The 139H2 in hybridoma culture supernatant was a kind gift from John Hilkins from The Netherlands Cancer Institute (NKI). The 139H2 was purified with Protein G Sepharose 4 Fast Flow beads (Merck), washed with PBS, eluted with 0.2 mM Glycine-buffer pH 2.5, neutralized with 1 M Tris-HCL pH 8 and dialyzed against PBS with Pierce Protein Concentrators PES, 30 kDa MWCO.

### Bottom-up proteomics – in-solution digestion

139H2 was denatured in 2% sodium deoxycholate (SDC), 200 mM Tris-HCl, and 10 mM Tris(2-carboxyethyl)phosphine (TCEP), pH 8.0 at 95 °C for 10 min, followed by 30 min incubation at 37 °C for reduction. The samples were then alkylated by adding iodoacetic acid to a final concentration of 40 mM and incubated in the dark at room temperature for 45 min. 3 μg sample was then digested by one of the following proteases: trypsin (Promega) and elastase (Sigma-Aldrich) in a 1:50 ratio (w/w) in a total volume of 100 μL of 50 mM ammonium bicarbonate at 37 °C for 4 h. After digestion, SDC was removed by adding 2 μL of formic acid (FA) and centrifuged at 14000× g for 20 min. Following centrifugation, the supernatant containing the peptides was collected for desalting on a 30 μm Oasis HLB 96-well plate (Waters). The Oasis HLB sorbent was activated with 100% acetonitrile and subsequently equilibrated with 10% formic acid in water. Next, peptides were bound to the sorbent, washed twice with 10% formic acid in water, and eluted with 100 μL of 50% acetonitrile/5% formic acid in water (v/v).

### Bottom-up proteomics – in-gel digestion

The hybridoma 139H2 was loaded on a 4%-12% Bis-Tris precast gel (Bio-Rad) in non-reducing conditions and run at 120 V in 3-Morpholinopropane-1-sulfonic acid (MOPS) buffer (Bio-Rad). Bands were visualized with Imperial Protein Stain (Thermo Fisher Scientific), and the size of the fragments evaluated by running a protein standard ladder (Bio-Rad). The Fab bands were cut and reduced by 10 mM TCEP at 37 °C, then alkylated in 40 mM IAA at RT in the dark, followed by alkylation in 40 mM IAA at RT in the dark. The Fab bands were digested by chymotrypsin and thermolysin at 37 °C overnight in 50 mM ammonium bicarbonate buffer. The peptides were extracted with two steps incubation at RT in 50% ACN, and 0.01% TFA, and then 100% ACN respectively.

### Bottom-up proteomics – LC-MS/MS

The peptides obtained by in-solution and in-gel digestion were vacuum-dried and reconstituted in 100 μL of 2% FA. The digested peptides were separated by online reversed-phase chromatography on an Agilent 1290 Ultra-high performance LC (UHPLC) or Dionex UltiMate 3000 (Thermo Fisher Scientific) coupled to a Thermo Scientific Orbitrap Fusion mass spectrometer. Peptides were separated using a Poroshell 120 EC-C18 2.7-Micron analytical column (ZORBAX Chromatographic Packing, Agilent) and a C18 PepMap 100 trap column (5 mm × 300, 5 μm, Thermo Fisher Scientific). Samples were eluted over a 90 min gradient from 0 to 35% acetonitrile at a flow rate of 0.3 μL/min. Peptides were analyzed with a resolution setting of 60 000 in MS1. MS1 scans were obtained with a standard automatic gain control (AGC) target, a maximum injection time of 50 ms, and a scan range of 350–2000. The precursors were selected with a 3 m/z window and fragmented by stepped high-energy collision dissociation (HCD) as well as electron-transfer high-energy collision dissociation (EThcD). The stepped HCD fragmentation included steps of 25, 35, and 50% normalized collision energies (NCE). EThcD fragmentation was performed with calibrated charge-dependent electron-transfer dissociation (ETD) parameters and 27% NCE supplemental activation. For both fragmentation types, MS2 scans were acquired at a 30 000 resolution, a 4e5 AGC target, a 250 ms maximum injection time, and a scan range of 120–3500.

### Bottom-up proteomics – peptide sequencing from MS/MS Spectra

MS/MS spectra were used to determine *de novo* peptide sequences using PEAKS Studio X (version 10.6)^39,40^. We used a tolerance of 20 ppm and 0.02 Da for MS1 and MS2, respectively. Carboxymethylation was set as fixed modification of cysteine and variable modification of peptide N-termini and lysine. Oxidation of methionine and tryptophan and pyroglutamic acid modification of N-terminal glutamic acid and glutamine were set as additional variable modifications. The CSV file containing all the *de novo* sequenced peptide was exported for further analysis.

### Bottom-up proteomics – template-based assembly via Stitch

Stitch (nightly version 1.4.0+802a5ba) was used for the template-based assembly^41^. The mouse antibody database from IMGT was used as template^42^. The cutoff score for the *de novo* sequenced peptide was set as 90 and the cutoff score for the template matching was set as 10. All the peptides supporting the sequences were examined manually.

### Cloning and Expression of recombinant 139H2 IgG and Fab

To recombinantly express full-length anti-MUC1 antibodies, the proteomic sequences of both the light and heavy chains were reverse-translated and codon-optimized for expression in human cells using the Thermo Fisher webtool (https://www.thermofisher.com/order/gene-design/index.html). For the linker and Fc region of the heavy chain, the standard mouse Ig γ-1 (IGHG1) amino acid sequence (Uniprot P01868.1) was used. An N-terminal secretion signal peptide derived from human IgG light chain (MEAPAQLLFLLLLWLPDTTG) was added to the N-termini of both heavy and light chains. BamHI and NotI restriction sites were added to the 5′ and 3′ ends of the coding regions, respectively. Only for the light chain, a double stop codon was introduced at the 3′ site before the NotI restriction site. The coding regions were subcloned using BamHI and NotI restriction-ligation into a pRK5 expression vector with a C-terminal octahistidine tag between the NotI site and a double stop codon 3′ of the insert, so that only the heavy chain has a C-terminal AAAHHHHHHHH sequence for nickel-affinity purification (the triple alanine resulting from the NotI site). After the sequence was validated by Sanger Sequencing, the HC/LC were mixed in a 1:1 DNA ratio and expressed in HEK293 cells by the ImmunoPrecise Antibodies (Europe) B.V company. After expression the culture supernatant of the cells was harvested and purified using a prepacked HisTrap excel column (Cytiva), following standard protocols. (see Supplementary Figure S2))

To recombinantly express anti-MUC1 Fab the coding regions of HC variable region were subcloned using AgeI and NheI restriction-ligation into a pRK5 expression vector. The subcloned region contains the mouse Ig γ-1 (IGHG1) Fab constant region with a C-terminal octahistidine tag followed by a double stop codon 3′ of the insert, so that only the heavy chain has a C-terminal AAAHHHHHHHH sequence for nickel-affinity purification (the triple alanine resulting from the NotI site). After the sequence was validated by Sanger Sequencing the HC/LC were mixed in a 1:1 (*m*/*m*) DNA ratio and expressed in HEK293 cells by the ImmunoPrecise Antibodies (Europe) B.V company. After expression the culture supernatant was loaded onto a 5 ml HisTrap excel column (Cytiva) using peristatic pump. Column was reconnected to the ÄktaGo system (Cytiva) for column wash (50 mM Tris at pH=8, 150 mM NaCl) and step elution (50 mM Tris at pH=8, 150 mM NaCl, 300 mM imidazole). Fraction from the peak corresponding to the Fab were concentrated using Amicon Ultra-15 (Millipore) and further purified by size-exclusion chromatography using Superdex 200 Increase 10/300 GL (Cytiva) in buffer 50 mM Tris (pH=8), 150 mM NaCl.

### Mammalian cell lines and culture conditions

The human gastrointestinal epithelial cell lines HT29-MTX^43^ and HT29-MTX ΔMUC1^5^ were cultured in 25 cm^2^ flasks in Dulbecco’s modified Eagle’s medium (DMEM) containing 10% fetal calf serum (FCS) at 37 °C in 10% CO^2^.

### Western blot

HT29-MTX and HT29-MTX ΔMUC1 lysates were prepared form cells grown to full confluency for 7 days in a 6-well plate. Cells were harvested by scraping and lysed with lysis buffer (10% SDS in PBS with 1× Halt Protease Inhibitor Cocktail). Concentration was measured by BCA-assay, 5× Laemmli buffer was added and sample was boiled for 15 min at 95 °C. A mucin-SDS gel was made according to Li et al.^5^; 40 μg of protein was added to each well and run in Boric acid-Tris buffer (192 mM Boric acid, Merck; 1 mM EDTA, Merck; 0.1% SDS, to pH 7.6 with Tris) at 25 mA for 1.5 h. Proteins were transferred to a polyvinylidene fluoride (PVDF) membrane using wet transfer for 3 h at 90 V/4 °C in transfer buffer (25 mM Tris; 192 mM glycine, Merck; 20% methanol, Merck). Afterwards, membranes were blocked with 5% BSA in TSMT (20 mM Tris; 150 mM NaCl, Merck; 1 mM CaCl_2_ (Sigma); 2 mM MgCl_2_, Merck; adjusted to pH 7 with HCl; 0.1% Tween 20 (Sigma)) overnight at 4 °C. The following day, membranes were washed with TSMT and incubated with 139H2 Wildtype, Synthetic or FAB antibodies (1:1000) in TSMT containing 1% BSA for 1h at RT. Membranes were washed again with TSMT and incubated with α-mouse IgG secondary antibody (A2304, Sigma) diluted 1:8000 in TSMT with 1% BSA for 1 h at RT, washed with TSMT followed by TSM. For detection of actin, cell lysates were loaded onto a 10% SDS-PAGE gel, transferred to PVDF membranes and incubated with α-Actin antibody (1:2,000; bs-0061R, Bioss) and α-rabbit IgG (1: 10,000; A4914, Sigma). Blots were developed with the Clarity Western ECL kit (Bio-Rad) and imaged in a Gel-Doc system (Bio-Rad).

### Western blot of MUC1 reporter constructs

Four MUC1 reporter constructs, expressed in engineered HEK293 cells, were a kind gift from Chistian Büll of the Copenhagen Center for Glycomics. Each reporter construct in 1× PBS was boiled in 5× laemmli buffer. 10 ng/25 ng of each construct was loaded per well on a 10% bis-acrylamide SDS gel for the 139H2/6× His-tag blots respectively. Samples were run in 1× Novex Tris-Glycine SDS Running Buffer (Thermo Fisher Scientific) for 1.5 h at 120 V. Proteins were transferred to a 0.2 μm Trans-Blot PVDF membrane (Bio-Rad) and transferred at 1.3 A/25 V for 7 min using the Trans-Blot Turbo system (Bio-Rad). Afterwards, membranes were blocked with 5% BSA in TSMT (20 mM Tris; 150 mM NaCl, Merck; 1 mM CaCl_2_, Sigma; 2 mM MgCl_2_, Merck; adjusted to pH 7 with HCl; 0.1% Tween 20, Sigma) overnight at 4 °C. The following day, membranes were washed with TSMT and incubated with 139H2 Wiltype, Synthetic antibody (1:1000) or HisProbe-HRP Conjugate (15165,Thermo Fisher Scientific,1:5000) in TSMT containing 1% BSA for 1h at RT. The 6× His-tag blots were washed with TMST and TSM and developed with the Clarity Western ECL kit (Bio-Rad) and imaged in a Gel-Doc system (Bio-Rad). The 139H2 membranes were washed again with TSMT and incubated with α-mouse IgG secondary antibody (A2304, Sigma) diluted 1:8000 in TSMT with 1% BSA for 1 h at RT, washed with TSMT followed by TSM and developed.

### Confocal microscopy

HT29-MTX and HT29-MTX ΔMUC1 cells were grown for 7 days to reach a confluent monolayer on cover slips (8 mm diameter#1.5) in 24-well plates. Cells were washed with Dulbecco’s Phosphate Buffered Saline (DPBS, D8537) and fixed with 4% paraformaldehyde in PBS (Affimetrix) for 30 min at RT. Fixation was stopped by adding 50 mM NH_4_Cl in PBS for 15 min. Cells were washed 2 times and permeabilized in binding buffer (0.1% saponin (Sigma) and 0.2% BSA (Sigma) in DPBS) for 30 min. Coverslips were incubated with 139H2 Wildtype, Synthetic of FAB at 1:100 dilution for 1h, washed 3× with binding buffer, incubated with Alexa Fluor-488-conjugated α-mouse IgG secondary antibodies (1:200; A11029, ThermoFisher) and DAPI at 2 μg/ml (D21490, Invitrogen) for 1 h. Coverslips were washed 3× with DPBS, desalted in MiliQ, dried and embedded in Prolong Diamond mounting solution (ThermoFisher) and allowed to harden. Images were collected on a Leica SPE-II confocal microscope with a 63× objective (NA 1.3, HCX PLANAPO oil). Controlled by Leica LAS AF software with default settings to detect DAPI, Alexa488, Alexa568 and Alexa647. Axial series were collected with step sizes of 0.29 μm.

### Surface Plasmon Resonance

N-terminally biotinylated synthetic MUC1 peptide with the sequence biotin-GGS-APDTRPAPG was ordered from Genscript. This was dissolved in PBS and printed on a planar streptavidin-coated SPR chip (P-Strep, SSens B.V.) using a continuous flow microfluidics spotter (Wasatch), flowing for 1 hour at RT, after which it was washed with SPR buffer (150 mM NaCl, 25 mM 4-(2-hydroxyethyl)-1-piperazineethanesulphonic acid (HEPES) with 0.005% Tween 20) for 15 min and quenched with biotin solution (10 mM biotin in SPR buffer). SPR experiments were performed using an IBIS-MX96 system (IBIS technologies) with SPR buffer as the running buffer. Dilution series of 2× steps of the full recombinant 139H2 or Fab were prepared, starting from a 10.0 μM stock for full IgG and a 7.88 μM stock for the Fab, diluting with SPR buffer. 20 dilution steps (including the stock) were used for the full IgG, and 10 dilutions were used for the Fab. SPR experiments were performed as a kinetic titration without regenerating in between association/dissociation cycles, with 30 min association and 10 min dissociation time for the full IgG and 6 min association and 4 min dissociation for the Fab. Binding affinity was determined by fitting data at binding equilibrium to a 2-site binding model for the full IgG and a 1-site (Langmuir) binding model for the Fab, using Scrubber 2.0 (Biologic software) and Graphpad Prism 5 (Graphpad software, Inc.).

### Crystallization and data collection

Sitting-drop vapor diffusion crystallization trials were set up at 20 °C by mixing 150 nl of complex with 150 nl of reservoir solution. The complex sample consisted of purified 139H2 Fab and MUC1 epitope peptide (APDTRPAPG; GeneScript) in a 1:2.5 molar ratio, at a total concentration of 3.8 mg/mL in a buffer of 50 mM trisaminomethane at pH 8.0 and 150 mM NaCl. The diffracting crystals grew in a condition of 0.2 M NaCl, 0.1 M sodium phosphocitrate, and 20% w/v Polyethylene glycol (PEG) 8000 used as reservoir solution. A 3:1 mixture of reservoir solution and glycerol was added as cryo-protectant to the crystals before plunge freezing them in liquid nitrogen. Datasets were collected at 100 K at Diamond Light Source beamline I24, equipped with an Eiger 9M detector (Dectris), at a wavelength of 0.6199 Å.

### Structure determination and refinement

Collected datasets were integrated using the xia2.multiplex pipeline^44^, and the three best datasets were subsequently merged and scaled in AIMLESS to a maximum resolution of 2.5 Å. Resolution limit cut off was determined based on mean intensity correlation coefficient of half-data sets, CC_1/2_. An initial model of 139H2 Fab was generated using ColabFold^45^. The variable region and constant region were placed in subsequent PHASER^46^ runs, the short linkers between the two regions were built manually and the CDRs were adjusted in COOT^47^. Clear density for the MUC1 peptide was present in the Fo-Fc map, and the peptide was built manually in COOT. The structure was refined by iterative rounds of manual model building in COOT and refinement in REFMAC5^48^. The final model was assessed using MolProbity^49^. All programs were used as implemented in CCP4i2 version 1.1.0^50^.

## Acknowledgements

This research was funded by the Dutch Research Council NWO Gravitation 2013 BOO, Institute for Chemical Immunology (ICI; 024.002.009) to J.S. KS and KCAPG are supported by the European Research Council under the European Union’s Horizon 2020 research and innovation program (ERC-2019-STG-852452). The authors would like to thank Diamond Light Source for beamtime (proposal mx25413), and the staff of beamline I24 for assistance with data collection.

## Data Availability

The raw LC-MS/MS files and analyses have been deposited to the ProteomeXchange Consortium via the PRIDE partner repository with the dataset identifier PXD043489. Coordinates and structure factors for 139H2 bound to the MUC1 epitope peptide have been deposited to the Protein Data Bank with accession code 8P6I.

