## Supplementary Information for "Reverse engineering the anti-MUC1 hybridoma antibody 139H2 by mass spectrometry-based *de novo* sequencing"

>139H2 Heavy Chain

QVQLQQSGAELVKPGASVKLSCKASGYTFTNYYMYWVLQRPGQGLEWIGEINPSNGGTTFNE
KFKNKATLTVDKSSSTAYMQLNSLTSEDSAVYYCTRSRYGNYVNYGMDYWGQGTSVTVSSAS
TTPPSVYPLAPGSAAQTNSMVTLGCLVKGYFPEPVTVTWNSGSLSSGVHTFPAVLQSDLYTLS
SSVTVPSSPRPSETVTCNVAHPASSTKVDKKIVPRDCGCKPCICTVPEVSSVFIFPPKPKDVLTI
TLTPKVTCVVVDISKDDPEVQFSWFVDDVEVHTAQTQPREEQFNSTFRSVSELPIMHQDWLNG
KEFKCRVNSAAFPAPIEKTISKTKGRPKAPQVYTIPPPKEQMAKDKVSLTCMITDFFPEDITVEW
QWNGQPAENYKNTQPIMNTNGSYFVYSKLNVQKSNWEAGNTFTCSVLHEGLHNHHTEKSLS
HSPGK

>139H2 Light Chain

DIVMSQSPSSLAVSVGEKVTMSCKSSQSLLYSNTQKNYLAWYQQKPGQSPKLLIYWASTRES
GVPDRFTGSGSGTDFTLTISSVKAENLAVYYCQQYYRYPPTFGGGTKLEIRRADAAPTVSIFPP
SSEQLTSGGASVVCFLNNFYPKDINVKWKIDGSERQNGVLNSWTDQDSKDSTYSMSSTLTLTK
DEYERHNSYTCEATHKTSTSPIVKSFNRNEC

*Supplementary Table S1*. Details of X-ray data collection, processing, and structure refinement.

| PDB accession code | 8P6I |
|---|---|
| **Data collection and processing** | |
| Space group | *C*2 |
| a, b, c (Å) | 211.8, 42.8, 129.1 |
| α, β, γ (deg) | 90.0, 122.3, 90.0 |
| Wavelength (Å) | 0.6199 |
| Resolution (Å) | 56.05-2.50 (2.60-2.50)* |
| Rmerge | 0.387 (1.813) |
| Rpim | 0.093 (0.673) |
| CC(1/2) | 0.988 (0.689) |
| No. of observations | 630424 (58514) |
| No. unique | 34554 (3836) |
| Mean I/σ (I) | 9.0 (0.7) |
| Completeness (%) | 100.0 (100.0) |
| Redundancy | 18.2 (15.2) |
| **Structure Refinement** | |
| Rwork/Rfree | 0.202/0.253 |
| Model composition | |
| Non-hydrogen atoms | 6719 |
| Protein residues | 854 |
| Water | 229 |
| B factors ($Å^2$) | |
| Protein | 33.6 |
| R.m.s. deviations | |
| Bond lengths (Å) | 0.0086 |
| Bond angles (°) | 1.53 |
| Validation | |
| MolProbity score | 1.53 |
| Clash score | 2.27 |
| Poor rotamers (%) | 3.12 |
| Ramachandran plot | |
| Favoured (%) | 97.19 |
| Allowed (%) | 2.69 |
| Outliers (%) | 0.12 |

*Values in parenthesis are for the highest resolution shell.

*Supplementary Table S2*. Overview of 139H2-MUC1 epitope contacts observed in crystal structure.

| **MUC1 peptide** | **group** | **139H2** | **group** | **interaction** |
|---|---|---|---|---|
| Ala1 | backbone | Arg99\|LC | sidechain | hydrogen bond |
| Pro2 | sidechain | Tyr98\|LC | sidechain | stacking |
| Asp3 | backbone | Tyr98\|LC | backbone | hydrogen bond |
| | sidechain | Arg99\|HC | sidechain | salt bridge |
| | sidechain | Tyr100\|LC | backbone | hydrogen bond |
| Thr4 | - | - | - | - |
| Arg5 | backbone | Tyr101\|HC | backbone | hydrogen bond |
| | backbone | Tyr102\|HC | sidechain | hydrogen bond |
| | sidechain | Glu50\|HC | sidechain | salt bridge |
| | sidechain | Thr59\|HC | sidechain | hydrogen bond |
| | sidechain | Tyr100\|LC | sidechain | stacking |
| Pro6 | backbone | Tyr33\|HC | sidechain | hydrogen bond |
| Ala7 | - | - | - | - |
| Pro8 | backbone | Ser54\|HC | sidechain | hydrogen bond |
| Gly9 | - | - | - | - |

*Supplementary Table S3.* Overview of reported MUC1-Fab structures. Structures in yellow contain O-glycopeptide, green unglycosylated peptide. The final column highlights the bound peptides in the context of MUC1's repeat region.

| **PDB ID** | **mAb** | **peptide** | **remarks** | **ref** | **VNTR** (2x repeat GSTAPPAHGVTSAPDTRPAP) |
|---|---|---|---|---|---|
| 7VAZ | 14A | RPAPGS(GalNAc)TAPPAHG | higher affinity for glycopeptide | https://doi.org/10.1101/2022.07.24.501275 | GSTAPPAHGVTSAPDT**RPAPGSTAPPAHG**VTSAPDTRPAP |
| 7V8Q | 14A | RPAPGST(GalNAc)APPAHG | higher affinity for glycopeptide | https://doi.org/10.1101/2022.07.24.501275 | GSTAPPAHGVTSAPDT**RPAPGSTAPPAHG**VTSAPDTRPAP |
| 7VAC | 14A | RPAPGS(GalNAc)T(GalNAc)APPAHG | higher affinity for glycopeptide | https://doi.org/10.1101/2022.07.24.501275 | GSTAPPAHGVTSAPDT**RPAPGSTAPPAHG**VTSAPDTRPAP |
| 7V4W | 16A | RPAPGSTAPPAHG | higher affinity for glycopeptide | https://doi.org/10.1101/2022.07.24.501275 | GSTAPPAHGVTSAPDT**RPAPGSTAPPAHG**VTSAPDTRPAP |
| 7V64 | 16A | RPAPGST(GalNAc)APPAHG | higher affinity for glycopeptide | https://doi.org/10.1101/2022.07.24.501275 | GSTAPPAHGVTSAPDT**RPAPGSTAPPAHG**VTSAPDTRPAP |
| 7V7K | 16A | RPAPGS(GalNAc)T(GalNAc)APPAHG | higher affinity for glycopeptide | https://doi.org/10.1101/2022.07.24.501275 | GSTAPPAHGVTSAPDT**RPAPGSTAPPAHG**VTSAPDTRPAP |
| 6TNP | 5E5 | APGST(GalNAc)AP | higher affinity for glycopeptide | https://doi.org/10.1039/D0CC06349E | GSTAPPAHGVTSAPDTRP**APGSTAP**PAHGVTSAPDTRPAP |
| 6KX1 | SN-101 | VTSAPDT(GalNAc)RPAPGSTA | higher affinity for glycopeptide | https://doi.org/10.1039/D0SC00317D | GSTAPPAHG**VTSAPDTRPAPGSTA**PPAHGVTSAPDTRPAP |
| 5T6P | AR20.5 | APDTRPAP | higher affinity for glycopeptide | https://doi.org/10.1093/glycob/cww131 | GSTAPPAHGVTS**APDTRPAP**GSTAPPAHGVTSAPDTRPAP |
| 5T78 | AR20.5 | APDT(GalNAc)RPAP | higher affinity for glycopeptide | https://doi.org/10.1093/glycob/cww131 | GSTAPPAHGVTS**APDTRPAP**GSTAPPAHGVTSAPDTRPAP |
| 5A2J | SM3 | APDTRP | higher affinity for glycopeptide | https://doi.org/10.1002/anie.201502813 | GSTAPPAHGVTS**APDTRP**APGSTAPPAHGVTSAPDTRPAP |
| 5A2K | SM3 | APDT(GalNAc)RP | higher affinity for glycopeptide | https://doi.org/10.1002/anie.201502813 | GSTAPPAHGVTS**APDTRP**APGSTAPPAHGVTSAPDTRPAP |
| 5A2I | SM3 | APDS(GalNAc)RP | higher affinity for glycopeptide | https://doi.org/10.1002/anie.201502813 | GSTAPPAHGVTS**APDTRP**APGSTAPPAHGVTSAPDTRPAP |
| 5A2L | SM3 | APDC(GalNAc)RP | higher affinity for glycopeptide | https://doi.org/10.1002/anie.201502813 | GSTAPPAHGVTS**APDTRP**APGSTAPPAHGVTSAPDTRPAP |
| 5N7B | SM3 | APDT(GalNAc)RP | higher affinity for glycopeptide; substitution in glycosidic linkage | https://doi.org/10.1021/jacs.8b13503 | GSTAPPAHGVTS**APDTRP**APGSTAPPAHGVTSAPDTRPAP |
| 6FRJ | SM3 | APDT(GalNAc)RP | higher affinity for glycopeptide; substitution in glycosidic linkage | https://doi.org/10.1021/jacs.8b13503 | GSTAPPAHGVTS**APDTRP**APGSTAPPAHGVTSAPDTRPAP |
| 6TGG | SM3 | APDT(GalNAc)RP | higher affinity for glycopeptide; with iminosugar | https://doi.org/10.1039/C9SC06334J | GSTAPPAHGVTS**APDTRP**APGSTAPPAHGVTSAPDTRPAP |
| 6FZR | SM3 | APDT(GalNAc)RP | higher affinity for glycopeptide; with fluorinated glycan | https://doi.org/10.1021/jacs.8b04801 | GSTAPPAHGVTS**APDTRP**APGSTAPPAHGVTSAPDTRPAP |
| 6FZQ | SM3 | APDT(GalNAc)RP | higher affinity for glycopeptide; with fluorinated glycan | https://doi.org/10.1021/jacs.8b04801 | GSTAPPAHGVTS**APDTRP**APGSTAPPAHGVTSAPDTRPAP |
| 5FXC | SM3 | APDT(GalNAc)RP | higher affinity for glycopeptide; glycan connected with linker | https://doi.org/10.1021/acs.joc.6b00833 | GSTAPPAHGVTS**APDTRP**APGSTAPPAHGVTSAPDTRPAP |
| 5OWP | SM3 | GVTSA(2fP)DT(GalNAc)RPAP | higher affinity for glycopeptide; with fluorinated proline | https://doi.org/10.1021/jacs.7b09447 | GSTAPPAH**GVTSAPDTRPAP**GSTAPPAHGVTSAPDTRPAP |
| 1SM3 | SM3 | TSAPDTRPAPGST | higher affinity for glycopeptide | https://doi.org/10.1006/jmbi.1998.2209 | GSTAPPAHGV**TSAPDTRPAPGST**APPAHGVTSAPDTRPAP |
| 8P6I | 139H2 | APDTRPAPG | | current work | GSTAPPAHGVTS**APDTRPAPG**STAPPAHGVTSAPDTRPAP |

Heavy chain

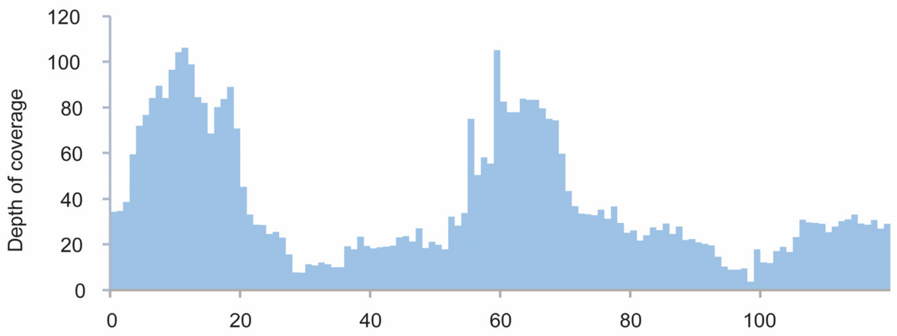


Light chain

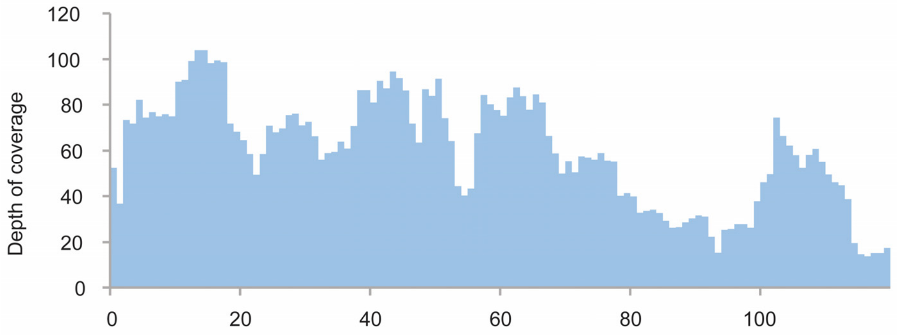


*Supplementary Figure S1.* Depth of coverage (total number of overlapping peptides mapped per position) for the variable domains of the 139H2 heavy and light chains.

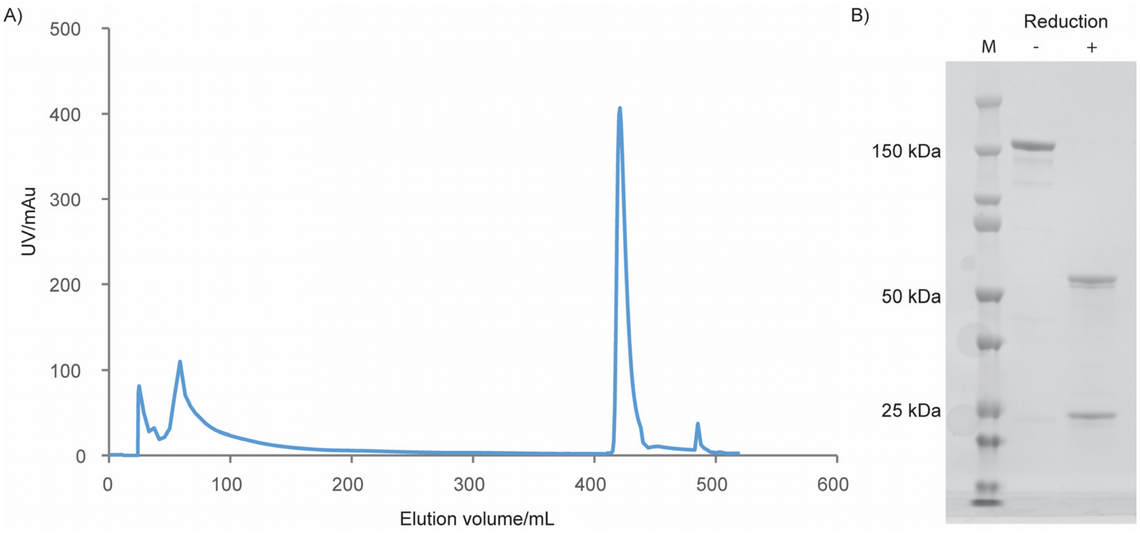


*Supplementary Figure S2.* Purification of recombinant 139H2. A) Elution profile across the imidazole gradient from the His-Trap purification. B) SDS-PAGE of the purified IgG product under reducing/non-reducing conditions.

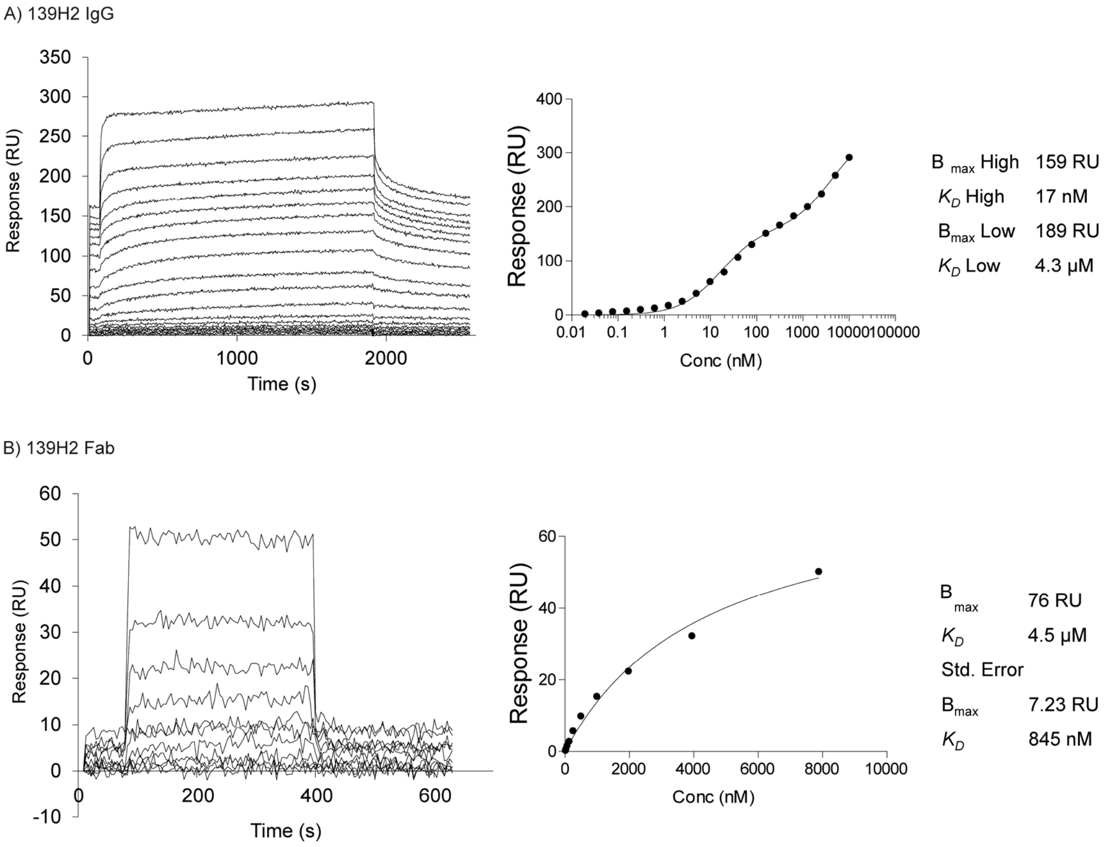


*Supplementary Figure S3*. Surface plasmon resonance quantification of binding affinity for the recombinant full-length 139H2 and its Fab to surface-immobilized MUC1 peptide. a) Binding of 139H2 full IgG to surface-immobilized biotinylated MUC1 peptide, analyzed by SPR. Equilibrium binding was fitted using a two-site binding model (right panel), with the respective $K_D$ and $B_{max}$ values for the high- and low-affinity binding indicated. b) Binding of 139H2 Fab to surface-immobilized biotinylated MUC1 peptide, analyzed by surface plasmon resonance SPR. Equilibrium binding was fitted using a one-site binding model (Langmuir isotherm, right panel), with $K_D$ and $B_{max}$ value indicated.

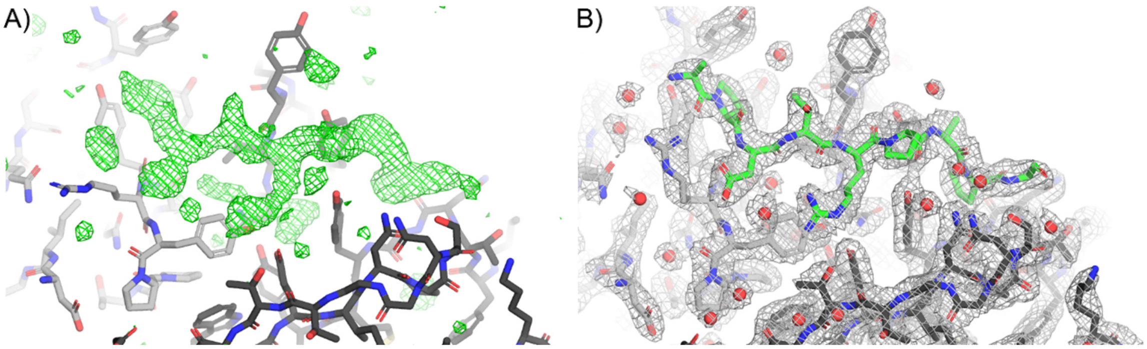


*Supplementary Figure S4.* Electron density for the MUC1 peptide bound to 139H2 Fab. (A) Positive $F_o$-$F_c$ omit density plotted at 3.0σ (green mesh), after correcting CDRs and refinement in REFMAC but excluding placement of the MUC1 peptide, shows well-resolved additional density at the peptide binding site. (B) The 2$F_o$-$F_c$ density plotted at 1.0σ (grey mesh) of the final refined model including waters (red spheres) shows a good fit for the modelled MUC1 peptide (green sticks). In both panels the 139H2 Fab molecule is shown in stick representation, with the light chain coloured in light grey, and the heavy chain coloured in dark grey, as in Figure 3.

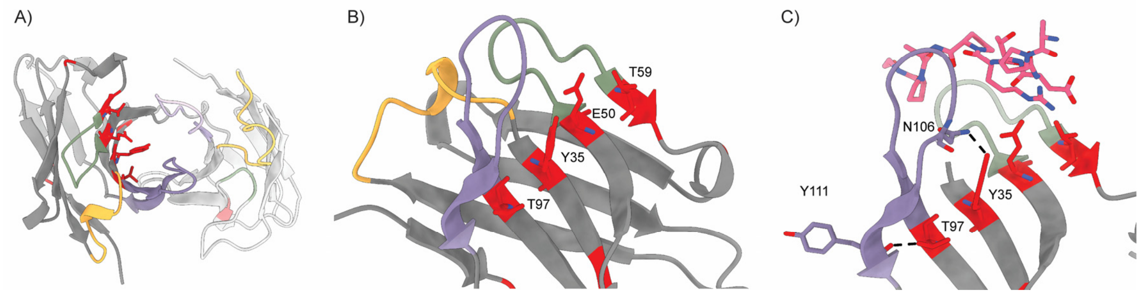


*Supplementary Figure S5.* Sites of somatic hypermutation in 139H2 framework. (A) Structure of 139H2 FAB with somatic hypermutations highlighted in red. (B) Hypermutations in heavy chain are organized in a stripe across the beta-sheet, side chains are oriented to the center of beta-barrel formed by heavy and light chains. (C) Interaction of Y35 and T97 with N106 and Y111, respectively, tilt CDR3 loop into the position where it can interact with MUC1.

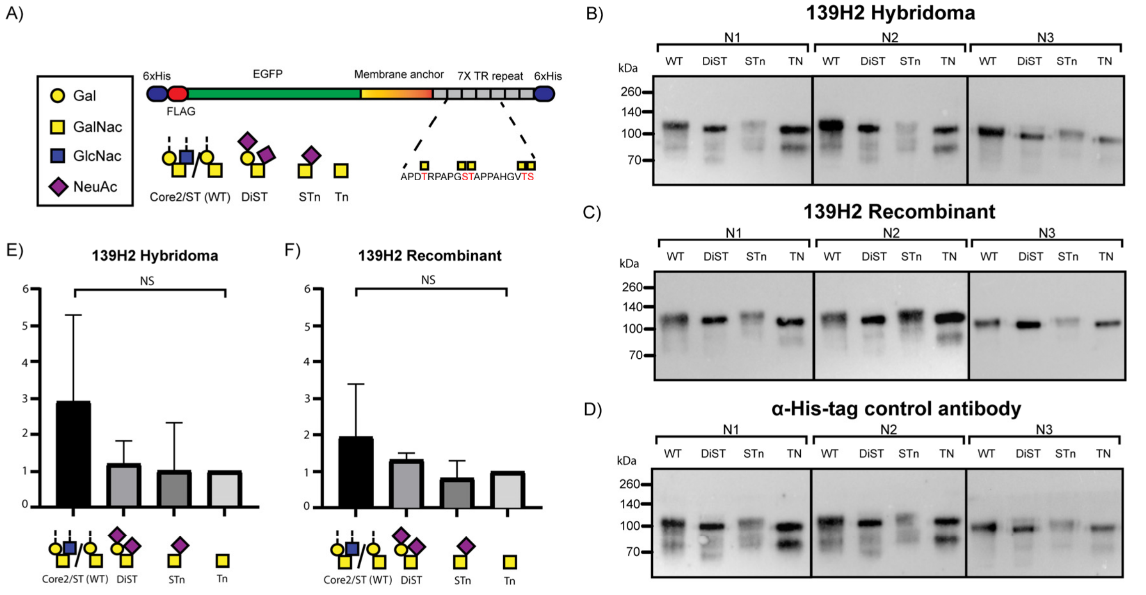


*Supplementary Figure S6.* Binding of 139H2 to MUC1 reporter constructs with different O-linked glycosylation. A) Schematic representation of the MUC1 Fragments used, adopted from Nason/Büll et al. 2021. The four fragments used contain 7 transmembrane repeats (TR) of MUC1 with 5 O-glycosylation sites with WT Core2/ST (WT)/ DiST/ STn or Tn glycan structures. Fragments were a kind gift from Christian Büll. Fig. 1B-D) Western blots against the MUC1 WT/DiST/STn/Tn fragments with 139H2 Hybridoma-derived antibody (B) (N=3), 139H2 Synthetic Recombinant antibody (C) (N=3) and a α-His-tag antibody control (D) (N=3). Fig. 1E) Western blot band intensities analyzed with Image Lab 6.0 software. Calculated intensity ratios were made relative to the intensity of MUC1-Tn. No significant difference in binding of 139H2 Hybridoma-derived or 139H2 Recombinant Synthetic was observed compared to the 6 α-His-tag antibody control.

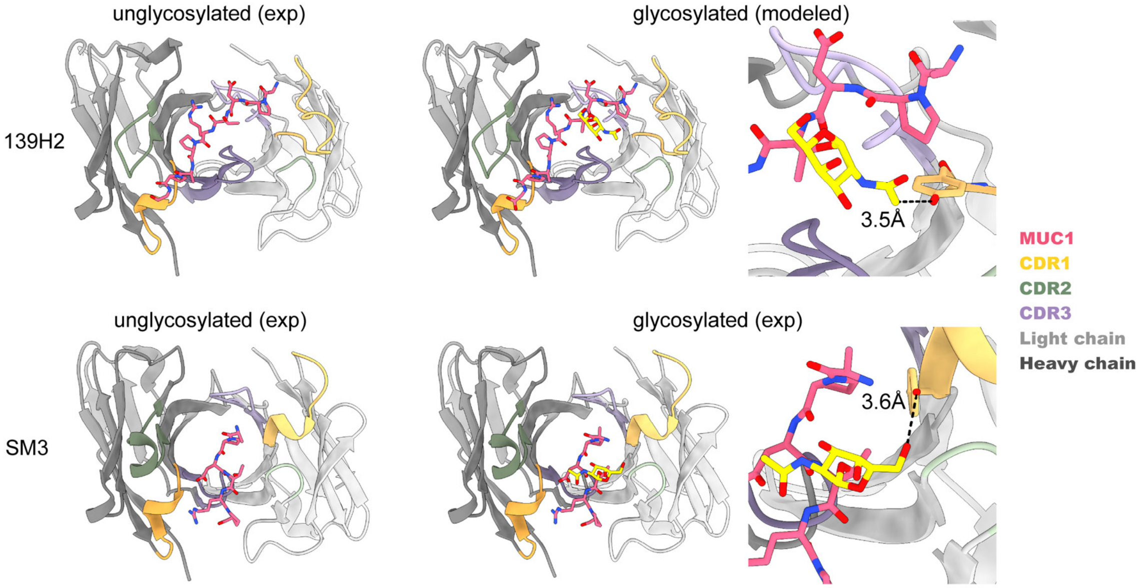


*Supplementary Figure S7*. Comparison of 139H2 and SM3 binding to MUC1. In SM3 the GalNAc residue makes an additional hydrogen bond with a tyrosine in CDRL1, similar interaction between T4 and GalNAc is predicted to be present also in 139H2.
